# Vulnerability of bats in Australia to wind turbine mortality: a trait-based assessment

**DOI:** 10.64898/2026.02.03.703634

**Authors:** Carissa V Harris, Christopher Turbill

## Abstract

Wind turbine collisions are a leading cause of bat mortality globally, yet the vulnerability of Australian bat taxa remains poorly understood. Trait-based assessments offer a valuable method for evaluating mortality risk in the absence of data on species- and region-specific turbine interactions. Behavioural and morphological traits are strongly associated with bat fatalities at wind farms globally. Here, we use a trait-based approach to identify the relative vulnerability of bat genera to the increasing threat of wind turbine fatalities in Australia. Through a global literature review, we identified three traits associated with increased mortality rates: foraging guild, movement distance, and wing morphology. For each trait we estimated the level of sensitivity at the genus level based on available published data. Sensitivity scores were summed to produce an overall vulnerability score. Our assessment identified *Saccolaimus* and *Taphozous* (Emballonuridae); *Austronomus, Chaereophon, Micronomus, Ozimops,* and *Setirostris* (Mollosidae); *Falsistrellus* (Vespertilionidae); and *Miniopterus* (Miniopteridae); as the most vulnerable genera. *Pteropus* (Pteropodidae) was assessed as high vulnerability for wing morphology and movement distance but low for foraging guild (as a non-aerial forager). However, because of their exceptional mobility, we assessed *Pteropus* as high overall vulnerability. We also attributed a confidence score to trait estimates, which reflected data availability. Confidence was rated as relatively high for foraging guild (61% genera rate as high) and wing morphology (50% high), and lowest for movement distance (14% high). Our framework provides a timely foundation for predicting mortality risk from the rapid expansion of wind energy infrastructure in Australia, which requires urgent conservation assessment and implementation of management actions to minimise the impacts. Our trait-based assessment also provides direction for targeted research to better understand and quantify the threat of wind turbine mortalities to populations of Australian bats.

## 1 Introduction

Renewable energy sources are becoming increasingly popular as an effective solution to mitigate the threat of climate change (GWEC 2025). Among these sources, wind energy is proposed as one of the most sustainable options, offering low cost and high energy yields (Veers et al. 2019). Driven by the adoption of the target set at the 28^th^ United Nations Climate Change Conference to triple renewable energy capacity by 2030, the wind energy industry is entering a phase of accelerated growth (GWEC 2025). However, the rapid global expansion of wind farms has brought some negative impacts such as noise pollution, visual disturbance, and threats to wildlife and their habitat (Saidur et al. 2011). Among wildlife, birds and bats are most affected (Schuster et al. 2015), with collisions with wind turbines identified as the leading cause of mass mortality events in bats globally (O’Shea et al. 2016).

Negative impacts of wind farms on bats include both direct mortalities and indirect effects that can also impact population growth (Schuster et al. 2015). Direct mortality typically results from collisions with moving turbine blades (Rydell et al. 2010). Indirect effects include habitat loss and displacement (Millon et al. 2018) and behavioural disruptions (Cryan et al. 2014). Any increase in mortality risk is especially concerning for bats because they generally have slow life histories (Barclay and Harder 2003), meaning their populations have a relatively low capacity to compensate by increased reproductive output (Purvis et al. 2000). Population sizes and trends of bats are poorly known, making it difficult to determine the extent of their response to declining numbers (Kunz et al. 2007). However, for one bat species in North America, models based on mortality data indicate the potential for turbine mortalities to cause substantial population-level declines (Frick et al. 2017). The uncertainty around how populations are impacted is concerning given that bats serve as valuable bioindicators of environmental health (Russo et al. 2021) and contribute importantly to ecosystem services that benefits agriculture production and public health (Kunz et al. 2011).

Australia is recognised as having some of the best wind resources in the world (GWEC 2025) and is part of a region that is experiencing rapid growth in renewable energy infrastructure (Xu et al. 2019). Yet, despite this potential for increased renewable energy, Australia and the surrounding region has the second lowest volume of published research on bat mortality linked to wind farms (Voigt et al. 2024). With the anticipated growth of wind energy in Australia, it is crucial to identify which bats are likely to face the highest risk of fatalities to mitigate this issue. While there is some monitoring of bat fatalities at wind farms most data are unpublished. Available mortality data is predominantly from southeastern Australia (Bennett et al. 2022; Hull and Cawthen 2013; Moloney et al. 2019; Smales 2012; Stark and Muir 2020). Using direct evidence to inform conservation efforts will lead to better outcomes, however, reliance on a currently small and localised set of mortality data to predict the vulnerability of all Australian bats increases the chances of a biased risk assessment (Christie et al. 2021).

While information on the interaction between bats and wind turbines is lacking for Australian species, there is a large amount of published data on mortality of bats at wind farms in other regions globally (Voigt et al. 2024). One method of assessing vulnerability to emerging anthropogenic threats, despite limited region-specific data, is by using traits known or suspected to be correlated with impacts (Gallagher et al. 2021). Traits can be defined as any measurable characteristic of an individual, and include morphological, phenological, physiological, or behavioural characteristics (Dawson et al. 2021). Classifying traits provides specific, comparable insights into how organisms interact with environmental pressures, and incorporating this information can improve the prioritisation and effectiveness of conservation actions (Gallagher et al. 2021; Green et al. 2022). By identifying which traits are associated with a particular threat, traits have been commonly used to assess vulnerability to climate change among a variety of taxa (Böhm et al. 2016; Conti et al. 2014; Foden et al. 2013) but have an under-appreciated potential for use in predicting outcomes of global change (Green et al. 2022).

Bat mortality at wind farms has been identified as being strongly associated with behavioural and morphological traits (Morant et al. 2025). Considering the availability of mortality data from other regions, and the lack of data availability in Australia, using traits to assess vulnerability to the imminent threat of wind farm mortality to Australian bats is a logical step forward. Our goal is to rate the relative vulnerability of Australian bats using a trait-based approach. To achieve this, we have three specific aims. First, we review the available literature and highlight traits related to the sensitivity and exposure of bats globally to wind farm fatalities. Second, we use these patterns to develop a trait-based assessment framework, identifying varying levels of sensitivity for each trait. Lastly, we apply this framework to Australian bats at the level of genus, providing a foundational ranking of vulnerability that can inform conservation assessment, support mitigation actions, and highlight research needs.

## 2 Methods

### 2.1 Trait selection

The selection of vulnerability-indicative traits was guided by reviewing global patterns of bat mortality at wind farms. To identify these patterns, a literature search was performed in Web of Science, Scopus, and Google Scholar using the search terms (bat* OR chiroptera) AND wind AND (farm* OR energy OR windfarm*) AND (mortality OR fatalities). Reference lists of included studies were also reviewed to identify additional relevant literature. From this process, we identified three traits consistently associated with increased mortality risk: foraging guild, movement distance, and wing morphology. We then structured our review around these traits, drawing from and highlighting key references that provide strong evidence for the inclusion of these traits in our assessment framework.

#### Foraging guild

Bats can be classified into guilds based on their foraging habitats and flight patterns, with open-space foragers typically flying high in open areas, edge-space foragers flying along habitat edges (e.g., forest-grassland borders), and narrow-space foragers flying within vegetation (Denzinger and Schnitzler 2013). Given these guilds provide a strong indicator of flight height, manoeuvrability, and habitat use, they have been used to predict the level of impact from wind farms (Ellerbrok et al. 2023; Morant et al. 2025; Roeleke et al. 2016). Open-space and edge-space foragers are considered at highest risk, because the height at which they move through space overlaps with the rotor-swept zone of wind turbines (Voigt et al. 2024). This correlation is confirmed when observing mortality patterns in North America, where the hoary bat (*Lasiurus cinereus*), eastern red bat (*Lasiurus borealis*), and silver-haired bat (*Lasionycteris noctivagans*) are most frequently recorded (Arnett et al. 2008). These species are known to forage above the canopy in open areas (Norberg and Rayner 1987). Similar patterns are documented in Europe (Brinkmann et al. 2006; Roeleke et al. 2016; Salguero et al. 2023), South Africa (Aronson 2022), and the Neotropics (Barros et al. 2015). Narrow-space foragers are considered low risk, however, may be impacted by habitat displacement (Ellerbrok et al. 2023; Voigt et al. 2024). Therefore, we scored foraging guild sensitivity as:

- **1 (low)** - Narrow-space foragers
- **2 (moderate)** - Edge-space foragers
- **3 (high)** - Open-space foragers

#### Movement distance

The movement distance of bats between roosting and foraging areas provides important insight into how and when they might be engaging with turbines (Cryan and Barclay 2009). Bats seem to commute and migrate along consistent routes that follow linear landscape features such as riparian vegetation, treelines, or coastlines (Ijäs et al. 2017; Pavey 1998; Roeleke et al. 2016). When turbines are placed along known routes, disproportionately higher mortality rates are recorded (Baerwald and Barclay 2009; Roeleke et al. 2016). In North America and Northwestern Europe, the bats most frequently reported are long-distance migratory species (Arnett et al. 2008; Rydell et al. 2010), although local species can also encounter wind turbines along their daily commuting routes (Roeleke et al. 2016). Overall, a global meta-analysis shows bat species that travel longer distances have greater collision rates than more sedentary species (Thaxter et al. 2017). Therefore, we scored movement distance sensitivity as:

- **(low)** - Short distance (<10 km)
- **(moderate)** - Moderate distance (10-100 km)
- **(high)** - Long distance (>100 km)

#### Wing morphology

Ecomorphology describes the causal relationship between an animal’s morphology and behaviour (Swartz et al. 2003). One of the most important ecomorphological traits in bats is wing morphology, which reflects flight performance across different habitat types (Denzinger and Schnitzler 2013). Since no single factor is likely to account for bat fatalities at wind farms across all species and regions, wing morphology serves as an integrative trait that captures a combination of factors (Crane et al. 2025). The common theme across mortality reports and reviews of impacts is bats that fly high in open space and commute long distances are most at risk (Voigt et al. 2024; Thaxter et al. 2017) and key parameters that influence these behaviours are wing loading and wing aspect ratio (Norberg and Rayner 1987).

Bats with higher wing loading (greater body mass relative to wing surface area) typically fly at higher speeds and cover longer distances (Norberg and Rayner 1987). Bats with higher aspect ratios (longer, narrower wings) experience less drag and improved aerodynamic efficiency during flight (Norberg and Rayner 1987). When considering foraging guilds, open-air foragers tend to have high aspect ratio wings because these wing shapes are more efficient for fast, sustained flight in open environments (Norberg and Rayner 1987). High aspect ratio and moderate wing loading is also seen in most long-distance migrants (Norberg and Rayner 1987). Narrow-space foraging bats typically have short, broad wings with low aspect ratios and low wing loading, which enhances manoeuvrability and enables slow, controlled flight in cluttered environments (Norberg and Rayner 1987). Bats that are more sedentary or engage in short-distance movements tend to have low aspect ratio and wing loading (Norberg and Rayner 1987) and are generally considered at lower risk (Voigt et al. 2024). We classified wing aspect ratio and wing loading following species comparisons presented by Norberg and Rayner (1987). Thresholds were based on distribution of measurements available for various bat species. Therefore, we scored wing morphology sensitivity as:

- **(low)** - Low aspect ratio & wing loading (<6)
- **(moderate)** - Moderate aspect ratio & wing loading (6-8)
- **(high)** - High aspect ratio & wing loading (>8)

### 2.2 Vulnerability framework

Trait-based vulnerability assessments often consider three factors: sensitivity, exposure, and adaptive capacity (Foden et al. 2013; Williams et al. 2008). However, modified approaches to this have been used when data is unavailable (Hyman et al. 2025).

Predicting exposure level to the threat of wind farm mortality is difficult. Specific locations of future wind infrastructure are often unknown, increasing the difficulty of anticipating where bats may encounter wind turbines. Additionally, there are still gaps in our knowledge about locations of bat populations, roosts, and migration or commuting routes. Instead, the traits we chose as indicators of sensitivity include elements of exposure. For example, bats capable of moving further distances are more likely to encounter turbines, and have increased exposure risk by indirect proxy (Thaxter et al. 2017). Exposure is also associated with habitat use (Williams et al. 2008), which is indicated in our assessment by foraging guild classification. For these reasons, we believe it is appropriate to omit exposure scores from our framework.

Adaptive capacity and resiliency serve as an indicator of whether a species can adapt to a stressor through physiological, behavioural and ecological changes, phenotypic plasticity or evolutionary responses, with high adaptive capacity often associated with traits characteristic of relatively fast life histories (Williams et al. 2008). The slow life histories of bats are associated with a lower adaptive capacity (Barclay and Harder 2003; Racey and Entwistle 2000). Bats have a limited ability to increase reproductive rate to counter reduced survival, and lower reproductive output constrains the ability to adapt to new stressors through genetic variation (Barclay and Harder 2003). Although population level responses have been modelled for one species (*L. cinereus*) in North America (Frick et al. 2017; Friedenberg and Frick 2021), we found no long-term studies that have identified the adaptive capacity of bats to the threat of wind farm mortality. Given the slow life-history traits of bats and the lack of empirical studies on their responses to wind farm mortalities, we consider adaptive capacity to be low across all bat genera. Our framework therefore defines vulnerability simply as the sum of the three traits associated with sensitivity (foraging guild, movement distance, and wing morphology), and the overall vulnerability score for each genus was calculated using the following formula:

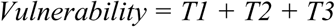

Where T1 is the assigned score for foraging guild, T2 for maximum movement distance, and T3 for wing morphology. The overall rating was then summed for a total vulnerability score of low: 0 to 4.9; moderate: 5.0 to 6.9; or high: 7.0 to 9.0.

### 2.3 Application of vulnerability framework

Rating of taxa was completed to genus level. Best estimates of each trait were derived through a literature search using combinations of keywords that included the relevant genus, or species name for members of the genus of interest. Given the recent taxonomic changes of Australian bats, each genus was screened for name changes and searched for accordingly. If further refinement was required, additional key words such as ‘foraging guild’, ‘movement’, ‘radio-telemetry’, ‘GPS’ or ‘wing morphology’ were included in the search.

Foraging guilds were assigned based on studies that directly observed flight behaviour (Barclay et al. 2000; Bullen and McKenzie 2001; McKenzie and Bullen 2009; McKenzie and Rolfe 1986; McKenzie et al. 2002; Knuckey et al. 2024; Pavey and Burwell 2004; Schulz 2000; Schulz and Hannah 1998; Tidemann et al. 1985; Williams and Thompson 2018), or Global Positioning System (GPS) tracking of fine scale movements (Gonsalves et al. 2024). Where direct observations are missing, guild was derived from the combination of echolocation characteristics and wing morphology (Adams et al. 2009; Denzinger and Schnitzler 2013; Mckenzie et al. 2020).

Movement distance was derived from maximum distance recorded during radio-tracking studies (Bondarenco et al. 2016; Bonaccorso et al. 2002; Gonsalves and Law 2017; Law 1993; Law and Anderson 2000; Law and Chidel 2004; Lumsden et al. 2002; McConville 2013; Pavey 1998; Schulz and Hannah 1998; Spencer and Fleming 1989; Winkelmann et al. 2003), banding (Dwyer 1966; Law et al. 2008) and GPS records (Bullen et al. 2023; Gonsalves et al. 2024; Tidemann and Nelson 2004). Where data were deficient, movement distance was inferred from wing morphology (Norberg and Rayner 1987; Mckenzie et al. 2020) or observational records (Bullen and McKenzie 2005).

Wing aspect ratio and wing loading were calculated as averages of measurements from several species within a genus (Bullen and McKenzie 2002; McKenzie et al. 2002; McKenzie et al. 2020; Norberg and Rayner 1987; Rhodes 2002; Schulz and Hannah 1998). When unavailable, wing morphology was derived from congeneric species from alternate regions using Norberg and Rayner (1987). Wing aspect ratio and wing loading were separately assigned a score from 1 to 3 based on the predefined criteria. The two scores were then combined, and the average was calculated for an overall wing morphology score for each genus.

### 2.4 Confidence of scoring

There are varying amounts of empirical and species-specific data available for Australian bats for the traits we have identified as predictive of sensitivity, therefore we also include a confidence score. Confidence scoring for each trait are assigned as follows:

#### Foraging guild

- **0 (low)** - Foraging guild not recorded, reliance on congeneric species
- **0.5 (moderate)** - Derived from echolocation characteristics and/or wing morphology
- **1 (high)** - Direct observation or GPS

#### Movement

- **0 (low)** - No movement records available for genus, reliance on congeneric species, inferred from wing morphology and/or echolocation characteristics, or presence/absence records
- **0.5 (moderate)** - Radio telemetry or banding recordings
- **1 (high)** - GPS

#### Wing morphology

- **0 (low)** - No wing morphology measurements available for any members of the genus, reliance on congeneric species
- **0.5 (moderate)** - Low (>10), or unknown sample size. Few members of the genus with measurements available
- **1 (high)** - Many available records of wing morphology. Many members of the genus with measurements available

The overall rating was then summed for a total confidence score of low: 0 to 1.0; moderate: 1.1 to 2.0; or high: 2.1 to 3.0.

## 3 Results

We assessed 28 Australian bat genera using a trait-based framework that includes three sensitivity traits: foraging guild, movement distance, and wing morphology. Each category was scored as low (1), medium (2), or high (3) risk (see Table 1 for references). Based on the cumulative trait vulnerability scores, 13 genera (46%) were considered low risk, 5 genera (18%) as moderate risk, and 10 genera (36%) as high risk (Table 1; Figure 1). The genera with the highest total vulnerability scores were *Falsistrellus*, *Miniopterus, Austronomus, Chaereophon, Micronomus, Ozimops, Setirostris, Saccolaimus, Taphozous,* and *Pteropus*. *Pteropus* was assessed as high vulnerability for movement distance and wing morphology, but low vulnerability for foraging guild due to classification as a narrow-space forager (Table 1). However, based on their exceptional mobility, we have assigned *Pteropus* as high risk for the purpose of this assessment. All members of the families Miniopteridae, Molossidae, and Emballonuridae were categorised as high risk, with *Austronomus, Saccolaimus,* and *Chaerephon* scoring high (3) for all trait categories (Table 1; Figure 1). Overall, 54% of genera had at least one trait assigned as high. High-risk genera typically accumulated a high score across all three trait categories, while low-risk genera were mostly composed of low trait scores (Figure 1). The most frequent trait increasing vulnerability was wing morphology, with 46% of taxa assigned a high score for this category (Figure 3). Foraging guild and movement showed equal contribution to the total scoring, with 25% of genera scored as high, 25% moderate, and 50% as low (Figure 3).

**Figure 1.**
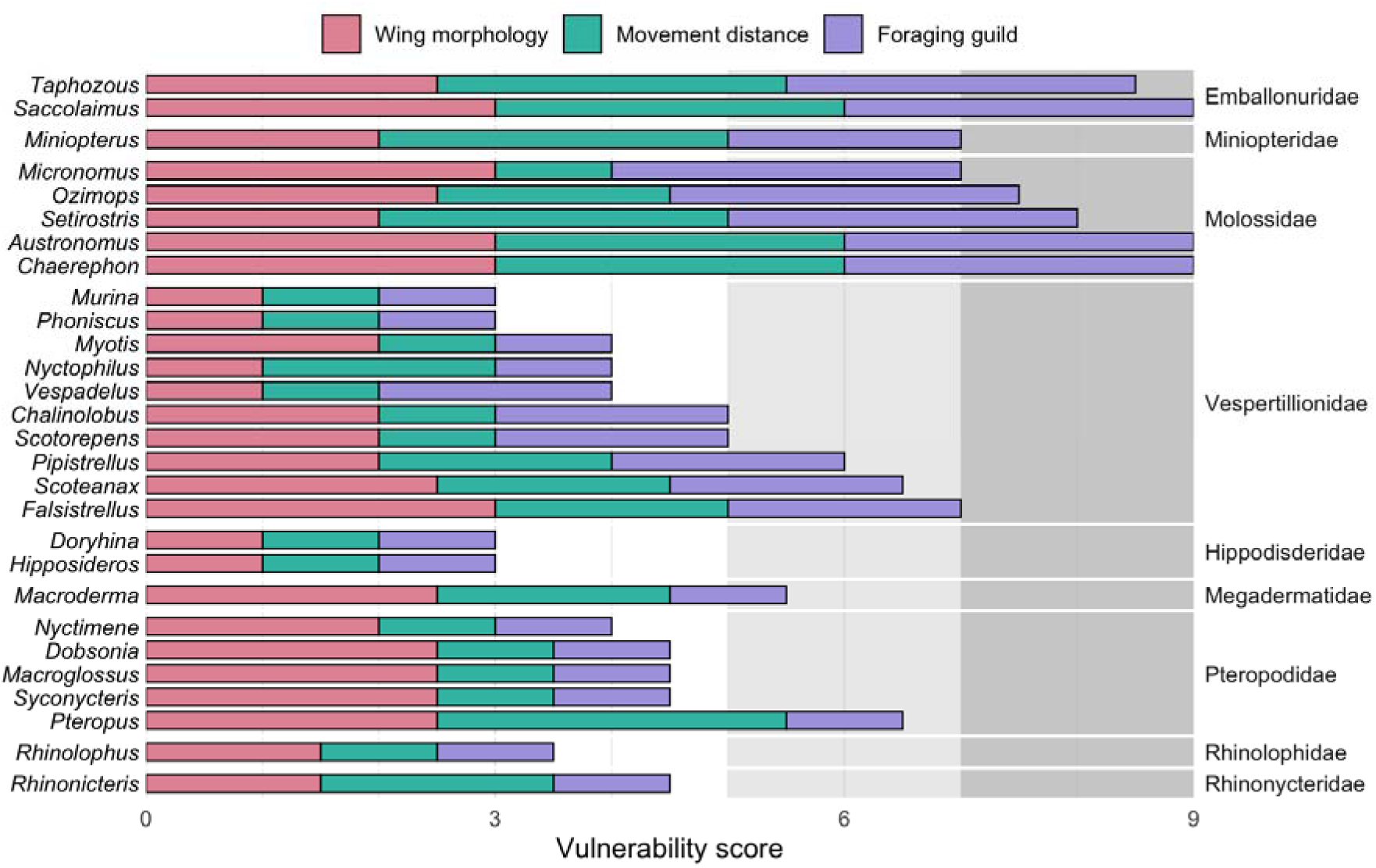
Vulnerability scores for genera of bats in Australia, which predict their risk of mortality from interaction with wind turbines. Vulnerability was assessed by assigning a score (1 to 3) for the traits of wing morphology (pink), movement distance (green) and foraging guild (purple), with the sum of scores representing total vulnerability. A score between 5 to 7 was considered a ‘moderate’ (light grey) and a score of 7 to 9 a ‘high’ (dark grey) level of vulnerability.

**Table 1.**
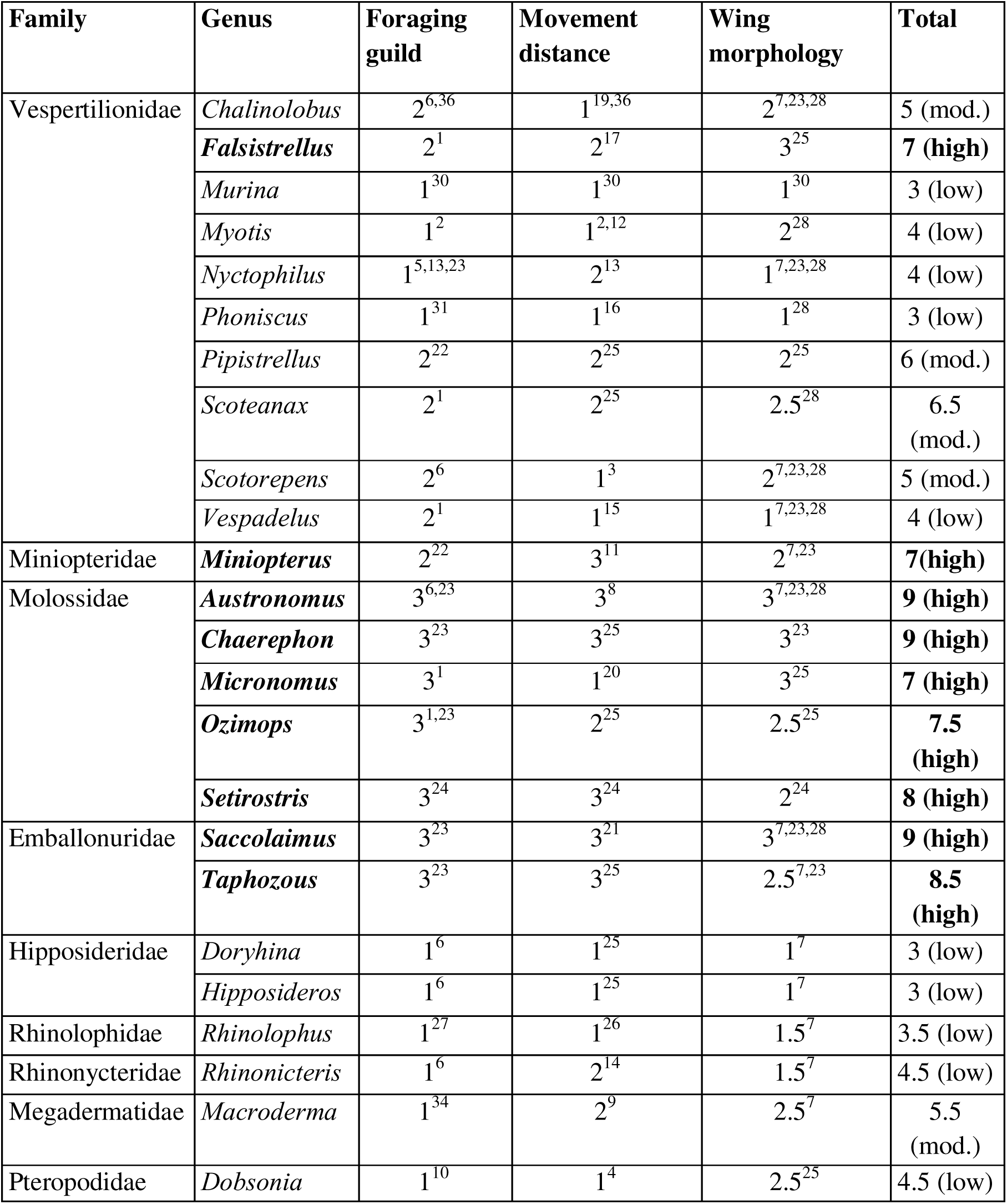
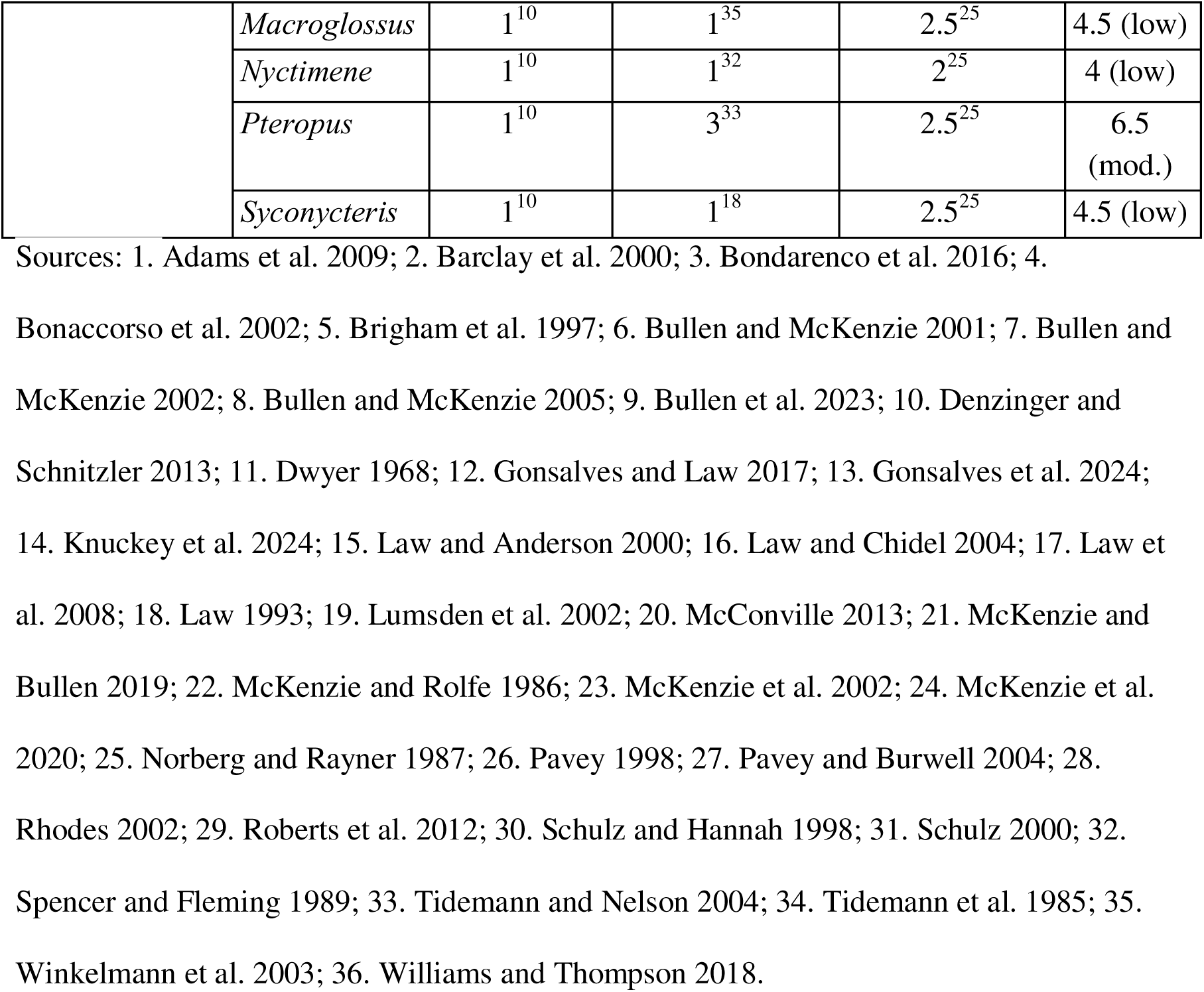
Scoring of traits included in the vulnerability assessment for genera of bats in Australia to the risk of mortality from interaction with wind turbines. Traits are combined to an overall vulnerability score.

Each trait was assigned a confidence score (Table 2). Of the 28 total confidence scores, 8 (29%) were considered high confidence, 15 (53%) moderate, and 5 (18%) low (Table 2; Figure 2). Overall, the genera with the highest confidence scoring were *Chalinolobus, Myotis, Nyctophilus, Phoniscus, Scotorepens, Miniopterus, Rhinolophus,* and *Macroderma,* while the lowest confidence were assigned to *Pipistrellus, Scoteanax, Setirostris, Hipposideros* and *Dobsonia* (Table 2; Figure 2). We had the greatest confidence in our scoring for foraging guild, with 61% of scores assigned as high (Figure 3). The lowest confidence was for our movement scores, with 39% assigned as low, 47% as moderate, and 14% as high (Figure 3).

**Figure 2.**
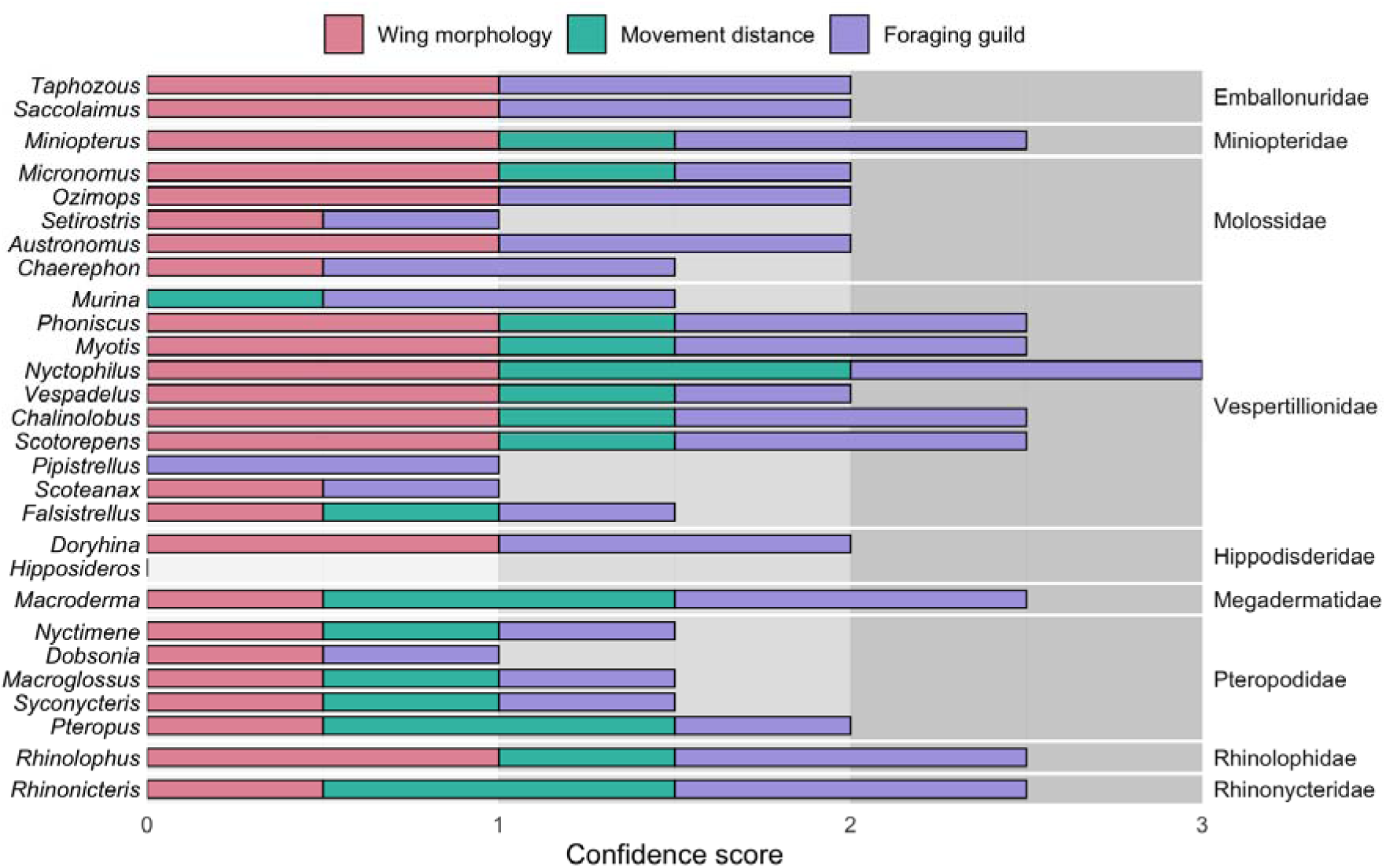
Confidence scores (0 = low, 0.5 = moderate, 1 = high) associated with each trait used to assess vulnerability for genera of bats in Australia.

**Figure 3.**
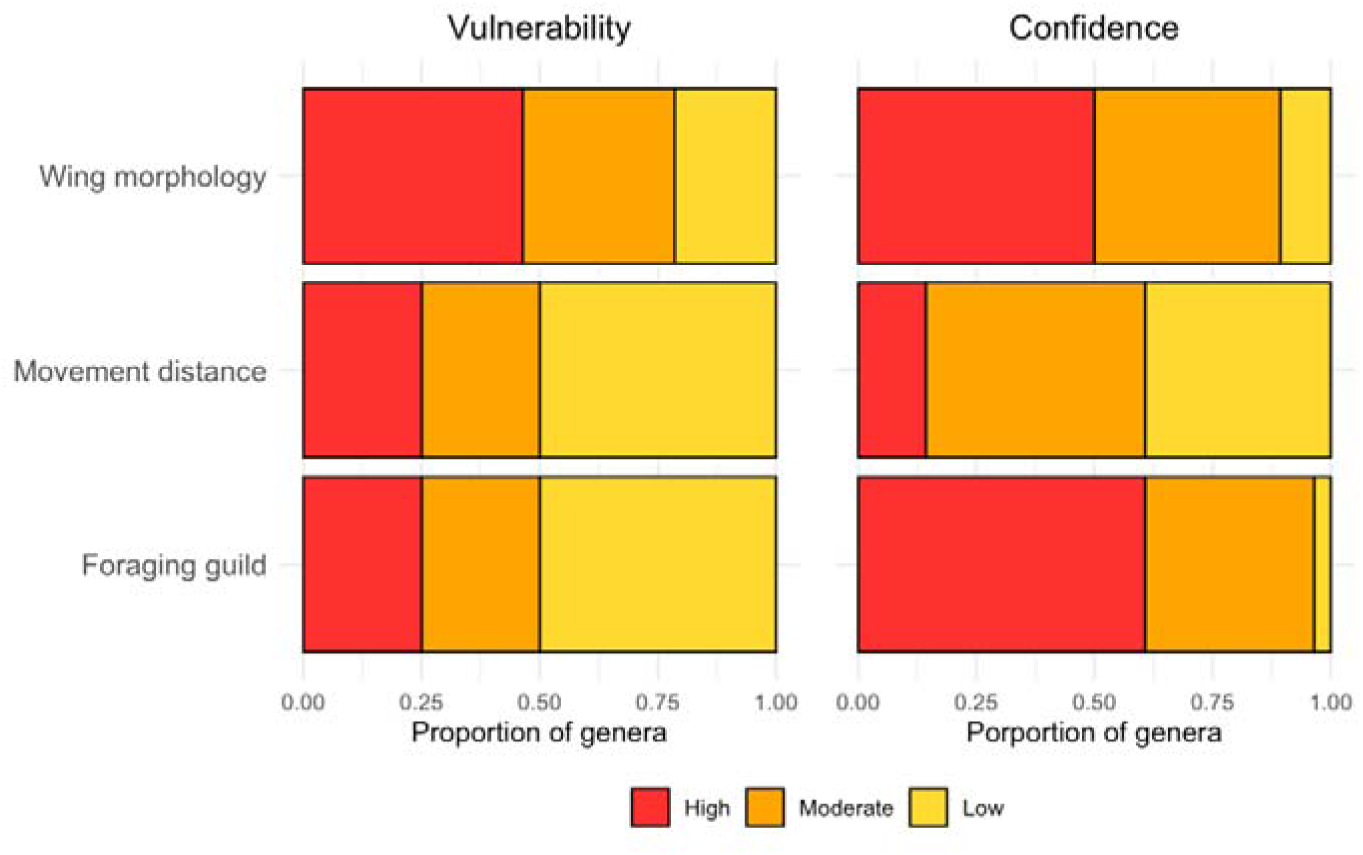
Proportion of the 28 genera of bats in Australia that were assigned low, moderate or high vulnerability and confidence scores for each trait.

**Table 2.**
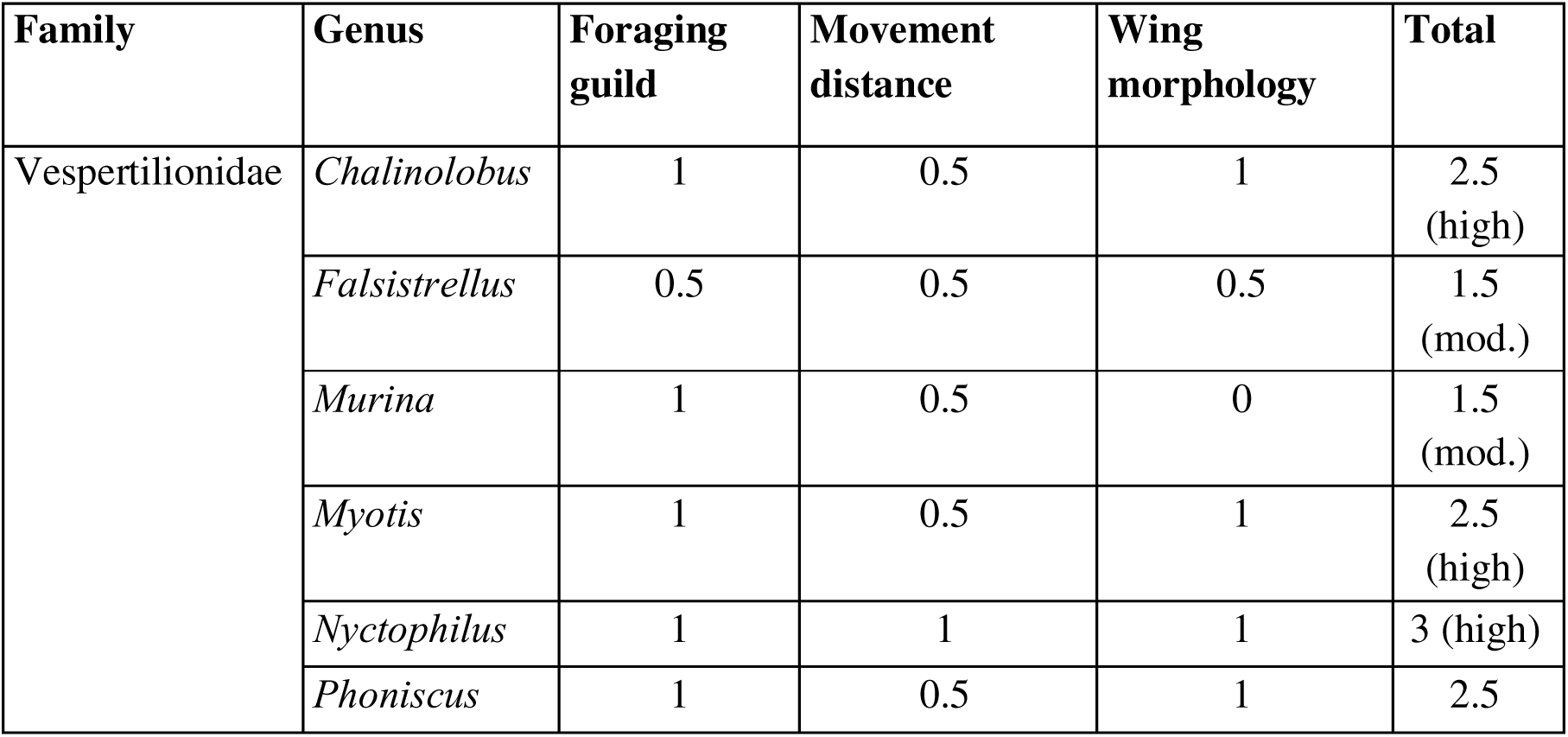
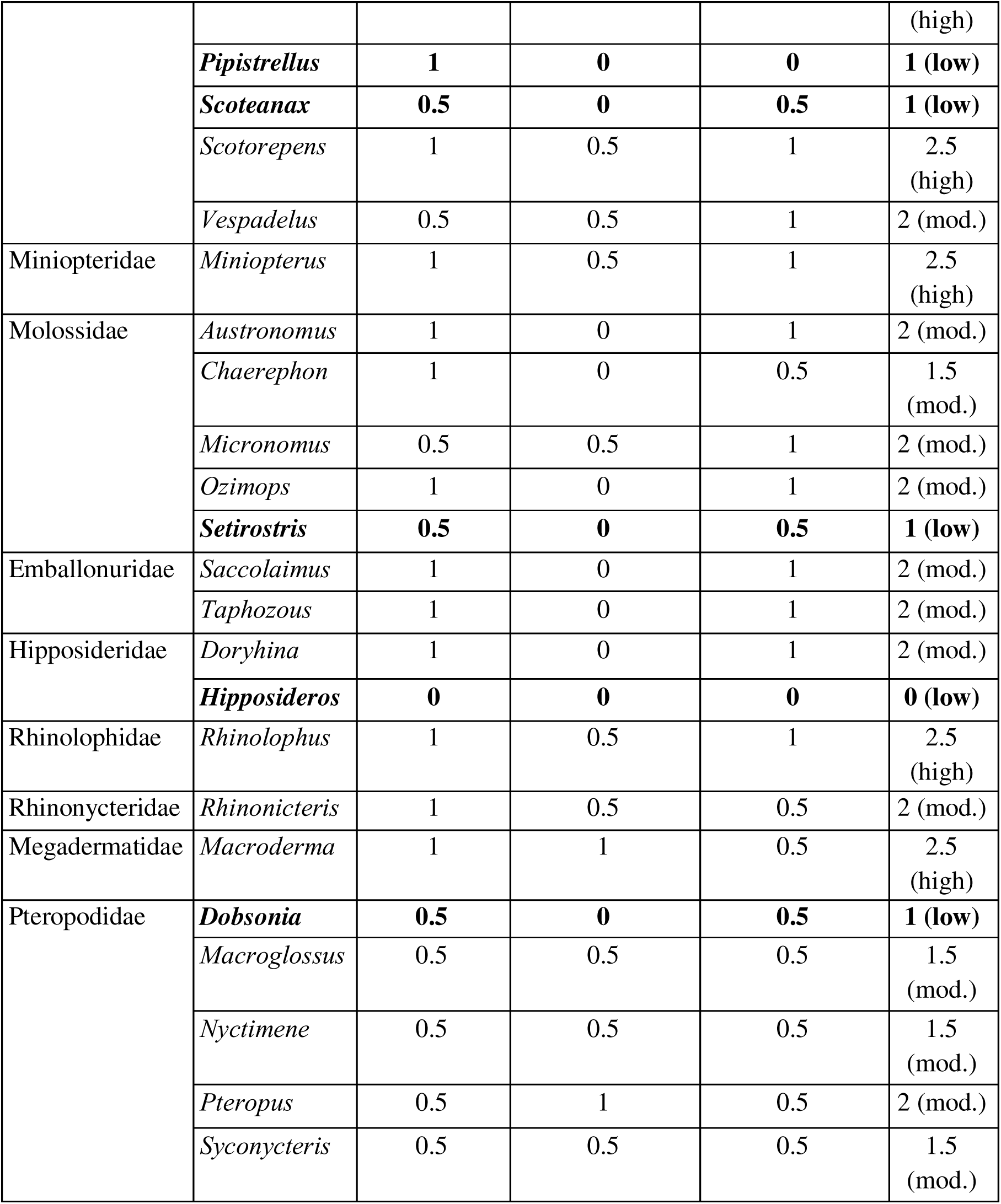
Confidence scores (0 = low, 0.5 = moderate, 1 = high) associated with each trait used to assess vulnerability for genera of bats in Australia.

## 4 Discussion

Globally, wind farms are a leading cause of bat mortality events (O’Shea et al. 2016). However, published research is concentrated in North America and Europe, leaving significant gaps in our understanding of impacts across other regions (Voigt et al. 2024). This raises concerns for how bats of varying geographic locations and phylogenetic diversity are being affected. Given this uncertainty, understanding region-specific vulnerability is a critical next step. Although we have limited knowledge on how Australian bats interact with wind turbines, conservation decisions often must be made despite large knowledge gaps (Frick et al. 2017). Failure to respond quickly to a rapidly declining population has already led to the extinction of an Australian bat, the Christmas Island pipistrelle *(Pipistrellus murrayi)*, which was a common and widespread species in its range (Martin et al. 2012). Taking a proactive approach to prevent taxa from future declines can reduce the likelihood that populations decline to critical levels, avoiding the high costs associated with species recovery plans (Walls 2018).

Our vulnerability assessment is consistent with published mortality data for Australian bats, providing some validation of its accuracy in predicting relative degree of threat. Our assessment scored the white-striped free-tailed bat (*Austronomus australis*) as high risk for all three traits. Mortality records of wind farms in Australia have found this species to be disproportionately impacted (Bennett et al. 2022; Hull and Cawthen 2013; Moloney et al. 2019; Smales 2012; Stark and Muir 2020). Members of the *Miniopterus* genus scored high overall vulnerability. Species belonging to this genus have recorded fatalities at wind farms in Australia (Bennett et al. 2022; Moloney et al. 2019; Stark and Muir 2020) including the critically endangered southern bent-wing bat (*Miniopterus orianae bassanii*). The high score for this genus is concerning given members are already facing a high risk of extinction. Other genera with recorded fatalities at wind farms in Australia which were assigned high scores using our assessment include *Falsistrellus* (Bennett et al. 2022; Moloney et al. 2019), *Micronomus* (Moloney et al. 2019), *Ozimops* (Moloney et al. 2019; Stark and Muir 2020), and *Pteropus* (Moloney et al. 2019; Stark and Muir 2020). Some genera (*Chaerophon, Sietrostris, Saccolaimus, Taphozous*) scored high, yet have no recorded fatalities in published reports. We do not believe this lessens the validity of the scores. Instead, the lack of empirical data on fatalities for these genera is likely caused by limited overlap between their distribution, the location of current wind energy infrastructure, and lack of centralisation and publication of mortality data. Therefore, we believe these genera are still highly vulnerable and have a greater risk of being impacted if wind farms are placed in their range.

While our framework assessed most genera with recorded fatalities as highly vulnerable, some were missed. For example, *Chalinolobus gouldii* has high mortality records at wind farms (Bennett et al. 2022; Hull and Cawthen 2013; Moloney et al. 2019; Smales 2012; Stark and Muir 2020), yet we scored members of the *Chalinolobus* genus as moderately vulnerable. Additionally, we scored genera *Nyctophilus* and *Vespadelus* as low risk however members of this genus have some recorded fatalities at wind farms (Bennett et al. 2022; Moloney et al. 2019; Smales 2012; Stark and Muir 2020).

### 4.1 Knowledge gaps and future research needs

Our assessment offers a foundation for understanding the relative vulnerability of bats in Australia to wind turbine mortalities, but it has limitations. While the ratings provided here are based on genera, we acknowledge that variation among species within genera is likely to exist for all three traits used. As a result, some genera could be given a less vulnerable score than expected for their species-specific traits. This is true particularly for more speciose genera. Members of the genus *Chalinolobus*, for example, show relatively large variation in aspects of their ecology. For the trait of wing morphology, *Chalinolobus gouldii* would be considered high risk, whereas *Chalinolobus dwyeri* would be moderate to low risk (Norberg and Rayner 1987). *Chalinolobus* species are broadly classified as edge-space foragers (Bullen and McKenzie 2001; Williams and Thomson 2018) however *Chalinolobus gouldii* has been observed foraging in open space (O’Neill and Taylor 1986). Our results provide only a relative measure of vulnerability among Australian genera and are not intended to be applied to all species within the genus. Where data allow, species-specific risks should be considered.

Foraging guilds are predictive of vulnerability however some species are variable in their foraging modes, making it difficult to classify them to a specific guild (Denzinger and Schnitzler 2013). This is the case for *Myotis* and *Macroderma*, two genera of bats in Australia that exhibit unique foraging styles (Barclay et al. 2000; Tidemann et al. 1985). For these genera, foraging guild alone is less likely to make them vulnerable. For example, placing wind farms near water sources may increase the vulnerability of *Myotis* substantially through habitat displacement (Scholz et al. 2025), and should be considered when determining risk level at wind farm sites.

Members of the family Pteropodidae typically fly close to vegetation during foraging and roosting, classifying them as narrow-space foragers (Denzinger and Schnitzler 2013). Following our framework, *Pteropus* species are considered to have moderate overall vulnerability to wind turbine collisions. However, due to their exceptional movement capabilities (Roberts et al. 2012; Welbergen et al. 2020), we have assessed them as high overall vulnerability. Vulnerability to wind turbine collision increases greatly with wide- ranging species (Thaxter et al. 2017). Therefore, using a precautionary approach, a high vulnerability score was deemed appropriate for this genus.

Movement distance knowledge is mainly restricted to home range foraging or commuting and does not account for dispersal and occasional long-distance movements. Overall, movement patterns of Australian bats are not well documented, with most of the information being derived from radio-tracking or predicted from wing morphology. While useful for understanding short range movements, radio-tracking has limitations (Harris et al. 1990). Radio-tracking is limited to tracking few individuals for a relatively short period of time; however individuals of different ages or sexes may exhibit different movement patterns over both space and time, and movements may occur rarely during migration events (Harris et al. 1990).

Studies analysing stable isotopes have been proposed as a promising technique for understanding the geographic origins and migration routes of bats (Cryan et al. 2014; Wright et al. 2020; Voigt et al. 2012). Stable isotopes in tissues of animals provide insight into where the tissue was grown (Seifert et al. 2018) although these methods provide less precision than GPS telemetry (Vander Zanden et al. 2018). Stable isotope analysis has proven useful in determining the origins of bats killed at wind turbine sites (Voigt et al. 2012; Baerwald et al. 2014; Pylant et al. 2016). Incorporating stable isotope analysis and GPS tracking to better understand movements of Australian bats is imperative for understanding movement patterns that increase vulnerability.

Movement behaviour is often influenced by temporal factors (Vardanis et al. 2011), however little is known about the seasonal movement patterns of Australian bat taxa. In North America, most mortality events of bats at wind turbines occur during late summer to early Autumn (Arnett et al. 2008; Thompson et al. 2017). This seasonal spike in mortality coincides with the migratory activity of bats most affected in this region (Thompson et al. 2017). A similar pattern has been observed in Europe (Brinkmann et al. 2006; Rydell et al. 2010; Georgiakakis et al. 2012). Because of this correlation, it is widely assumed that there is a causal link between seasonal migration and mortality events (Arnett et al. 2008; Kunz et al. 2007). Although distinct temporal patterns are observed in bat fatalities in the northern hemisphere (Arnett et al. 2008; Rydell et al. 2010), these patterns appear less pronounced in the southern hemisphere (Aronson 2022). Seasonal migratory patterns in tropical and subtropical regions are more likely shaped by alternate pressures such as resource availability rather than climatic constraints (Popa-Lisseanu and Voigt 2009). Additionally, breeding behaviours (e.g., swarming) occur at specific times of year and may influence when bats encounter turbines (Cryan and Barclay 2009). To inform effective mitigation strategies additional efforts should be directed towards understanding temporal movement and behavioural patterns of Australian bat taxa.

Wing morphology may offer insight into the vulnerability of species in specific regions based on the strong correlation between flight height and morphological traits. Wing morphology for Australian species is restricted to relatively few studies, and more efforts should be made to record wing aspect ratio and wing loading in Australian bats. Wider availability of these measurements would allow for stronger predictions of species-specific vulnerability. When measurements for genera are missing, foraging guild and movement capability are often inferred from wing morphology or echolocation characteristics. However, wing morphology does not always predict movement distance with perfect accuracy. A study that used radio-telemetry to track *C. gouldii* found both sexes travelled further (6.9km) between roost and foraging areas than wing morphology predicted (3.0km) (Lumsden 2002). While the use of wing morphology as a proxy for movement is a useful technique, further research confirming this correlation is required.

The goal of our work was to determine the relative vulnerability of Australian bats to the threat of wind turbine mortality. Our choice of framework was based on the urgency of assessing this risk and available knowledge from observing global trait patterns. Trait-based approaches for conservation are considered the best alternative to direct measurement when based on credible evidence (Gallagher et al. 2021). Such assessments are useful when extrapolating patterns across a range of ecosystems, regions, and taxonomic groups (Foden et al. 2013). We believe the abundance of research completed in alternate regions provides reliable indication of the traits associated with high risk to wind turbine collision. Despite the limitations of this assessment, our results are useful as a foundation for understanding which Australian bat genera have the highest relative mortality risk. We believe this assessment can be used to guide conservation priorities, research efforts, and mitigation strategies to reduce population declines caused by this emerging and potentially important threat.

